# *In Vitro* Activity of a Novel Metal-Based Antimicrobial against Multidrug-Resistant *Klebsiella pneumoniae*

**DOI:** 10.64898/2026.02.27.708516

**Authors:** Raquel L. Almeida, Nuno A. Faria, Mariana Araújo, Cláudia Malta-Luís, Francisco C. Mendes, Oscar Lenis-Rojas, Beatriz Royo, Maria Miragaia

## Abstract

Multidrug-resistant (MDR) *Klebsiella pneumoniae*, classified by the World Health Organization (WHO) as a critical priority pathogen, represents a global health thereat requiring novel antimicrobials urgently. Here we evaluated the *in vitro* antimicrobial activity of a novel iridium-based compound (OMKP-3), against MDR *K. pneumoniae*. OMKP-3 exhibited robust antimicrobial activity in M9 minimal media (MIC=6.25µg/mL) and rapid bactericidal effect (MBC=12.5µg/mL) against the tested MDR *K. pneumoniae* strains. OMKP-3 showed antibiofilm ability and was active against multiple MDR Gram-negative pathogens, including *Escherichia coli*, *Enterobacter cloacae*, *Pseudomonas aeruginosa* and *Serratia marcescens* (MIC range:6.25-25µg/mL). Importantly, OMKP-3 showed no cytotoxicity against mammalian cells after 24 hours of exposure. When combined with polymyxin B, OMKP-3 acted as an adjuvant, enhancing polymyxin B activity (FIC≤0.5). OMKP-3 was less prone to induce high-level resistance in MDR *K. pneumoniae* compared to ciprofloxacin, and supressed the growth of resistant bacteria at a low and non-cytotoxic concentration (4xMIC). *K. pneumoniae* strains harboring truncated Ompk35/36 porin genes exhibited higher OMKP-3 MICs, indicating that these porins may serve as an important entry pathway. Spectrometry analysis revealed that OMKP-3 was able to accumulate intracellularly (1.57µg/mL), with minimal Resistance-Nodulation-Division (RND) efflux pump extrusion involvement. Furthermore, analysis of the resistant mutant, harboring a mutation in the outer membrane protein DegS, together with fluorescence microscopy, suggests that OMKP-3 induces membrane-associated damage. No cross-resistance between OMKP-3 and commonly used antibiotics was observed. Collectively, these findings identify OMKP-3 as a promising novel antimicrobial agent against MDR *K. pneumoniae*, likely acting through an unexplored bacterial target.

**Importance:** Multidrug-resistant (MDR) *Klebsiella pneumoniae* is a critical global health threat and is among the leading causes of hospital0hyphenorendash;associated mortality, largely due to the scarcity of effective therapeutic options. Alarmingly, the current antimicrobial pipeline fails to address this issue, relying largely on derivatives of existing scaffolds that offer only short-term clinical benefit due to rapid resistance emergence. Developing antibiotics against Gram-negative pathogens is particularly challenging because of their highly impermeable outer membrane and efficient efflux systems, limiting intracellular drug accumulation. Metal-based antimicrobials emerge as a promising alternative. Our findings showed that OMKP-3, an iridium complex, exhibits potent bactericidal activity against MDR *K. pneumoniae* without selecting for high-level resistance, suggesting the potential for sustained therapeutic efficacy. Additionally, it demonstrated to accumulate intracellularly with minimal efflux involvement. Together, these features position OMKP-3 as a valuable and underexplored novel antimicrobial strategy for addressing the escalating threat of MDR *K. pneumoniae* infections.

## Introduction

*Klebsiella pneumoniae*, is a Gram-negative opportunistic pathogen that represents a serious health threat, particularly in hospital settings. It is a leading cause of healthcare-associated infections, including urinary tract infections, pneumonia and bacteremia (1, 2), and is associated to increased mortality and morbidity in immunocompromised patients, the elderly (1) and neonates (3). Besides nosocomial infections, *K. pneumoniae* can cause severe and invasive community-acquired infections in healthy individuals, particularly in Asian countries (4).

The treatment of *K. pneumoniae* infections relies on antibiotic therapy; however, the high levels of antimicrobial resistance (AMR) reported for this pathogen worldwide is of serious concern (5). In Europe (EU/EEA), resistance to third-generation cephalosporins remains the most frequently reported resistance phenotype in *K. pneumoniae*, reaching 32.9% in 2024, followed by fluoroquinolones (31.4%) and aminoglycosides (21.5%) (6). Of particular concern is the increasing trend in carbapenem resistance over the years, rising from 8.5% in 2017 (7) to 11.3% in 2024 (6). Because carbapenems are critical for the treatment of severe infections caused by Multidrug-resistant (MDR) *K. pneumoniae*, their diminished efficacy poses a serious clinical challenge (8).

In 2019, *K. pneumoniae* was responsible for approximately 193 000 deaths directly attributed to AMR worldwide, ranking as the third most common pathogen responsible for antimicrobial resistance (AMR)-related mortality (9). Strains resistant to carbapenems and third-generation cephalosporins were implicated in more than half of these fatalities (9), underscoring the absence of effective therapeutic alternatives to treat infections caused by these antibiotic-resistant strains.

As a Gram-negative bacteria, *K. pneumoniae* employs membrane-mediated resistance mechanisms that present an enormous challenge to antimicrobial design (10). These strategies include reduced antimicrobial uptake via porins - a major route of antibiotic entry (10) - and active extrusion of antimicrobials via efflux pumps (11), which altogether limit intracellular drug accumulation. The capacity of *K. pneumoniae* to form biofilms on host infections and in abiotic surfaces, such medical devices, further contributes to persistent infections and increased antibiotic tolerance (12). Collectively, these features emphasize the urgent need for novel antimicrobial agents capable of overcoming the complex resistance landscape of this pathogen.

To aggravate the scenario, it is unlikely that the current antimicrobial pipeline will adequately address this clinical need in the near future (13) . According to recent World Health Organization (WHO) reports, only four agents in the clinical pipeline target CR-Kp, all belonging to existing β-lactams/β-lactamase inhibitor classes (14). Additionally, over 80% of antibacterial agents approved between 2017 and 2023 derive from known chemical classes and lack new molecular targets or modes of action, rendering them vulnerable to cross resistance and rapid resistance emergence (14). Altogether, the high resistance and mortality levels, and dry antimicrobial pipeline, position antibiotic-resistant *K. pneumoniae* among the WHO critical priority pathogens for the development of new antibiotics (13). In response to this unmet need, non-traditional antimicrobial strategies have gained increasing attention. Metal-based complexes, also referred to as metalloantibiotics, have emerged as promising antimicrobial candidates in recent years. Their renewed prominence follows decades of application in anticancer therapy and their influential early use in combating bacterial infections in the 20th century (15). These compounds consist of a central metal ion coordinated to a diverse array of ligands, a feature that enables rational structural modification and the development of derivatives with optimized antimicrobial properties (15). Importantly, metal-based complexes can engage in unique mechanisms of action, including enzyme inhibition through substrate mimicry, generation of reactive oxygen species (ROS), and covalent binding to essential biomolecules (16). In a recent study by Frei and co-workers (17) which evaluated a large and chemically diverse compound library, metal-based complexes stood out for their high success rates compared to organic compounds, combining potent antimicrobial activity with low cytotoxicity. Additionally, a higher proportion of metal-based complexes were found to be active against Gram-negative bacteria, suggesting an enhanced ability to circumvent the impermeable outer membrane (17). Among studied metals, iridium complexes demonstrated particularly high success rates (17). Nevertheless, the development of iridium-based antimicrobials remains in its early stages, and their mode of action are still largely unexplored relative to other metal-based therapeutics.

To address this knowledge gap, we investigated the antimicrobial potential of an iridium-based compound against MDR *Klebsiella pneumoniae*. By merging antimicrobial susceptibility testing, cytotoxicity assessment, and exploratory mechanistic analyses, this work aims to evaluate the potential of iridium-based complexes as a foundation for the development of innovative antimicrobial strategies capable of overcoming the permeability and resistance barriers of Gram-negative bacteria.

## Results

### OMKP-3 chemical characteristics

OMKP-3 is a half-sandwich iridium complex featuring an N-heterocyclic carbene (NHC) ligand (18) (**Figure S1**). The complex has a molecular weight of ≈700 Da and exhibits moderate aqueous solubility (18 µM), partly attributable to the presence of amide functional groups within the ligand framework. Both the cyclopentadienyl moiety and the NHC ligand contribute to the structural and kinetic stability of the complex, while the chloride ligands provide labile coordination sites capable of undergoing substitution under physiological conditions, thereby facilitating potential interactions with biomolecules. OMKP-3 can be synthesized reproducibly in straightforward manner with an overall yield of 80% and at low production cost.

### OMKP-3 antimicrobial activity in different growth media

Metal-based antimicrobials agents have been described to exhibit lower antimicrobial activity when tested in complex bacterial growth media (19). Based on this evidence, OMKP-3 antimicrobial activity was assessed for their antimicrobial activity in both standard rich media (Mueller-Hinton broth) and minimal media (M9 minimal media) against MDR *K. pneumoniae* (S10p9A and OND234 strains) and antibiotic-susceptible *E. coli* (ATCC25922). Consistent with previous reported observations, OMKP-3 exhibited reduced activity in rich growth medium (minimum inhibitory concentration (MIC) >200µg/mL) compared to M9 minimal medium (MIC=6.25µg/mL) (**Table 1**). To further explore the association between higher antimicrobial activity and the growth media, we tested two hypothesis: i) interactions between metal-based complexes and components of the rich media may contribute to loss of antimicrobial activity, specially the presence of protein-based organic components (19); ii) growth in M9 medium may impose nutritional stress on *K. pneumoniae*, thereby increasing its susceptibility to additional stressors, including the action of antimicrobials.

**Table 1.**
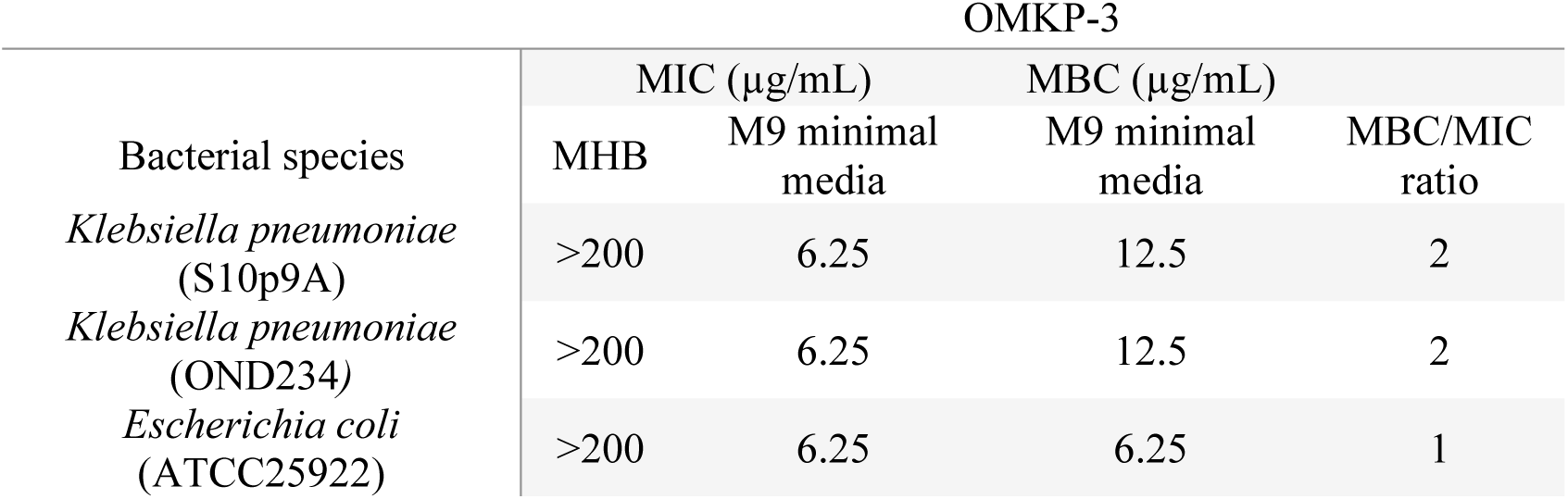
OMKP-3 minimum inhibitory concentrations (MICs) against Multidrug-resistant (MDR) *Klebsiella pneumoniae* (S10p9A and OND234 strains) and *Escherichia coli* (ATCC25922) in standard rich media (Muller-Hinton broth (MHB)) and M9 minimal media. Minimum bactericidal concentrations (MBC) and MHB/MIC ratio of OMKP-3 in M9 minimal medium against the same bacterial strains.

A great proportion of rich growth media composition, such as Muller-Hinton, is attributed to amino acids, components that are absent in M9 minimal media. Therefore, to explore the first hypothesis we have tested OMKP-3 antimicrobial activity in M9 minimal media supplemented with 0.0002%, 0.005%, 0.8% and 1.6% of amino acids from casein hydrolysate against *K. pneumoniae* OND234. Our results showed that while at lower amino acids concentrations (0.0002% and 0.005%) the OMKP-3 MIC remained unchanged, at higher concentrations (0.8% and 1.6%) OMKP-3 MIC was increased by >8-fold, suggesting that the existence of amino acids in the growth media is directly contributing to the loss of OMKP-3 activity.

To further test the second hypothesis, we have examined the activity of common antibiotics with distinct mechanisms of action (colistin, chloramphenicol, gentamicin, ciprofloxacin and trimethoprim) in the same nutrient limited conditions (M9 minimal media) against MDR *K. pneumoniae* (S10p9A and OND234) and *E. coli*. Contrary to OMKP-3, all tested antibiotics showed unchanged or decreased antimicrobial activity in minimal medium (≥ MICs), when compared to Mueller-Hinton broth (**Table 2**). This pattern suggests that the enhanced activity in M9 medium is a specific feature of OMKP-3 rather than a broad response to nutrient limitation.

**Table 2.** Minimum inhibitory concentrations (MICs) for common antibiotics (colistin, chloramphenicol, gentamicin, ciprofloxacin and trimethoprim) and OMKP-3 against Multidrug-resistant (MDR) *Klebsiella pneumoniae* and *Escherichia coli* using M9 minimal media and Muller-Hinton broth (MHB).

| Antibiotic | MIC (µg/mL) |  |  |  |  |  |
| --- | --- | --- | --- | --- | --- | --- |
|  | *K. pneumoniae* S10p9A |  | *K. pneumoniae* OND234 |  | *E. coli* ATCC25922 |  |
|  | MHB | M9 media | MHB | M9 media | MHB | M9 media |
| Colistin | 0.25 | 2 | 0.25 | 1 | <0.006 | <0.006 |
| Chloramphenicol | 32 | 32 | 2 | 4 | 2 | 2 |
| Gentamicin | 0.25 | 0.25 | 16 | 64 | 0.25 | 1 |
| Ciprofloxacin | 0.125 | 0.25 | 0.5 | 0.5 | <0.0004 | <0.0004 |
| Trimethoprim | 2 | 8 | 128 | 128 | 0.25 | 0.25 |
| OMKP-3 | >200 | 6.25 | >200 | 6.25 | >200 | 6.25 |

### OMKP-3 exhibited rapid bactericidal activity

OMKP-3 bactericidal activity was confirmed by observing a >3-log reduction in the number of colony forming units (CFUs) after 18±2h of complex exposure (minimal bactericidal activity (MBC) =12.5µg/mL for *K. pneumoniae*; MBC=6.25µg/mL for *E. coli*) (**Table 1**). Its bactericidal effect was further validated when we calculated OMKP-3 MBC/MIC ratios for both species. For tested *Enterobacteriaceae* MBC/MIC ratios were lower than 4 (*K. pneumoniae*: MBC/MIC=2; *E. coli*: MBC/MIC=1), indicating that the complex has potent bactericidal activity against these bacteria where bacterial killing occurred at concentrations similar to those of bacterial inhibition.

To further characterize OMKP-3 bactericidal activity, its kinetic of action was determined by time-kill kinetic assays. *K. pneumoniae* (S10p9A and OND234) and *E. coli* (ATCC25922) were exposed to different concentrations of OMKP-3 ranging from 1x to 8xMIC (6.25µg/mL to 50µg/mL) for 24 hours, and bacterial viability (CFU/mL) was assessed over time (**Figure 1**). For all tested bacteria and in the conditions examined, OMKP-3 exposure at the MIC (6.25µg/mL) was not sufficient to kill all bacterial cells in the population over 24 hours. At 2xMIC (12.5µg/mL), OMKP-3 exhibited bactericidal activity against MDR *K. pneumoniae* during the first 8 hours, yet there was bacterial regrowth afterwards. For *E. coli*, the same concentration of complex led to 100% bacterial killing after 4 hours of OMKP-3 exposure, indicating greater OMKP-3 susceptibility. At 4xMIC (25µg/mL), OMKP-3 was able to effectively kill *K. pneumoniae* strains after 4 hours of incubation and *E. coli* after 2 hours. At 8xMIC, OMKP-3 exhibited 100% bacterial killing after 2 hours, against the three tested bacterial strains. OMKP-3 demonstrated to have, generally, increasing antimicrobial activity with rising concentrations, characteristic of concentration-dependent antimicrobial agents.

**Figure 1.**
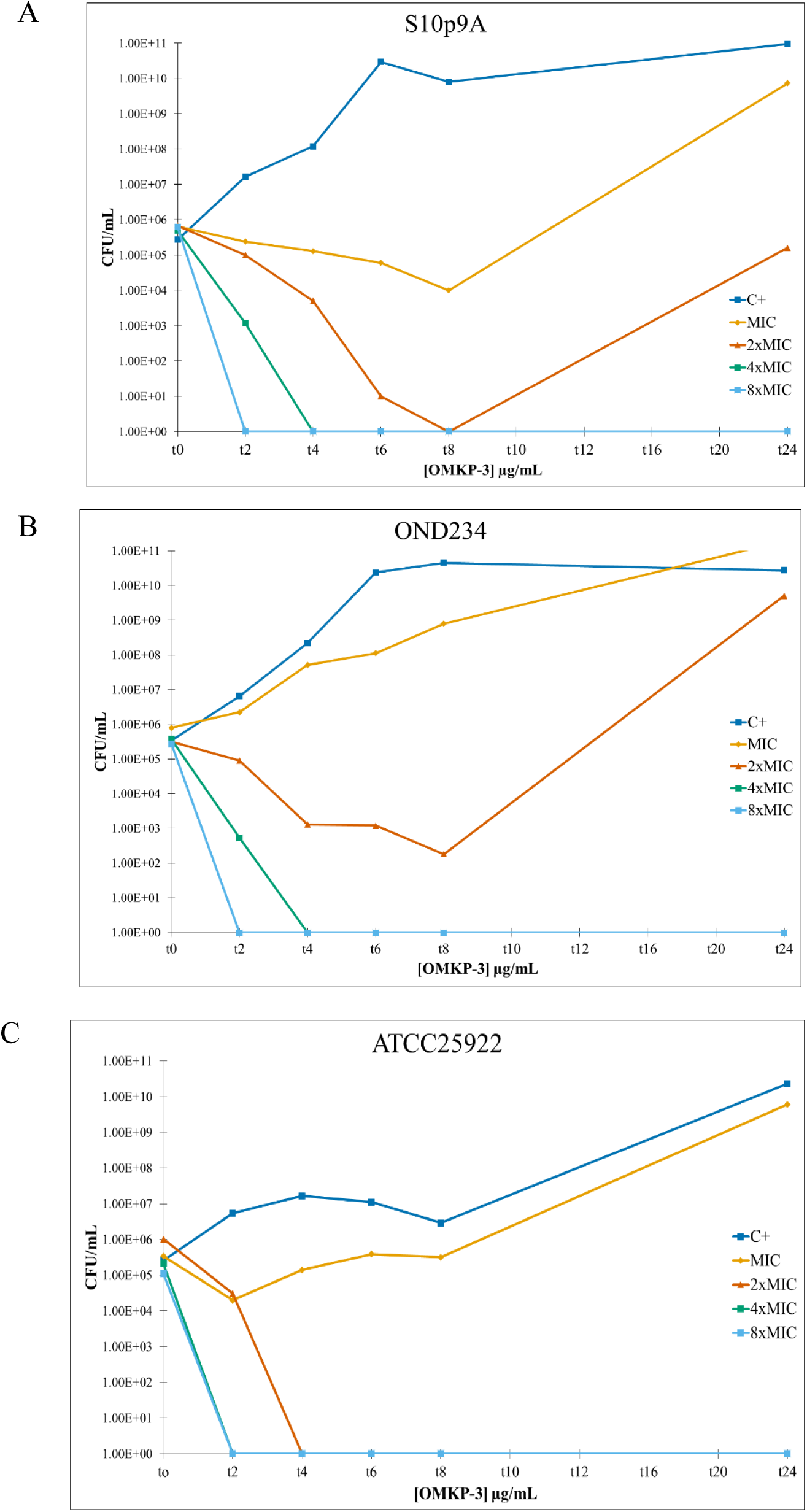
OMKP-3 time-kill curves against *Klebsiella pneumoniae* S10p9A (A) and OND234 (B) and *Escherichia coli* ATCC25922 (C) strains exposed to different OMKP-3 concentrations (MIC - 8xMIC) for 24h. Bacterial viability (CFU/mL) was monitored at selected timepoints.

### OMKP-3 was active against MDR *Klebsiella pneumoniae* and other MDR Gram-negative bacteria

Given OMKP-3 high antimicrobial potential against the tested MDR *K. pneumoniae* strains, we assessed OMKP-3 efficiency in the broader *K. pneumoniae* population, we tested its antimicrobial activity against additional 46 clinical carbapenem-resistant and MDR *K. pneumoniae* strains belonging to diverse genetic backgrounds, accounting for 23 sequence types (STs) (**Table S1**) (20). Moreover, we evaluated its spectrum of activity against other MDR Gram-negative pathogens.

Our results showed that OMKP-3 MIC varied between 6.25µg/mL and 50µg/mL in the *K. pneumoniae* population tested (**Figure 2**), with an estimated MIC50 and MIC90 of 25 µg/mL and 50µg/mL respectively. Importantly, OMKP-3 was also active against other clinically important MDR Gram-negative bacteria (*Enterobacter cloacae*, *Escherichia coli*, *Serratia marcescens* and *Pseudomonas aeruginosa*) ranging from 6.25µg/mL and 25µg/mL, showing broad-spectrum potential (**Table 3**).

**Figure 2.**
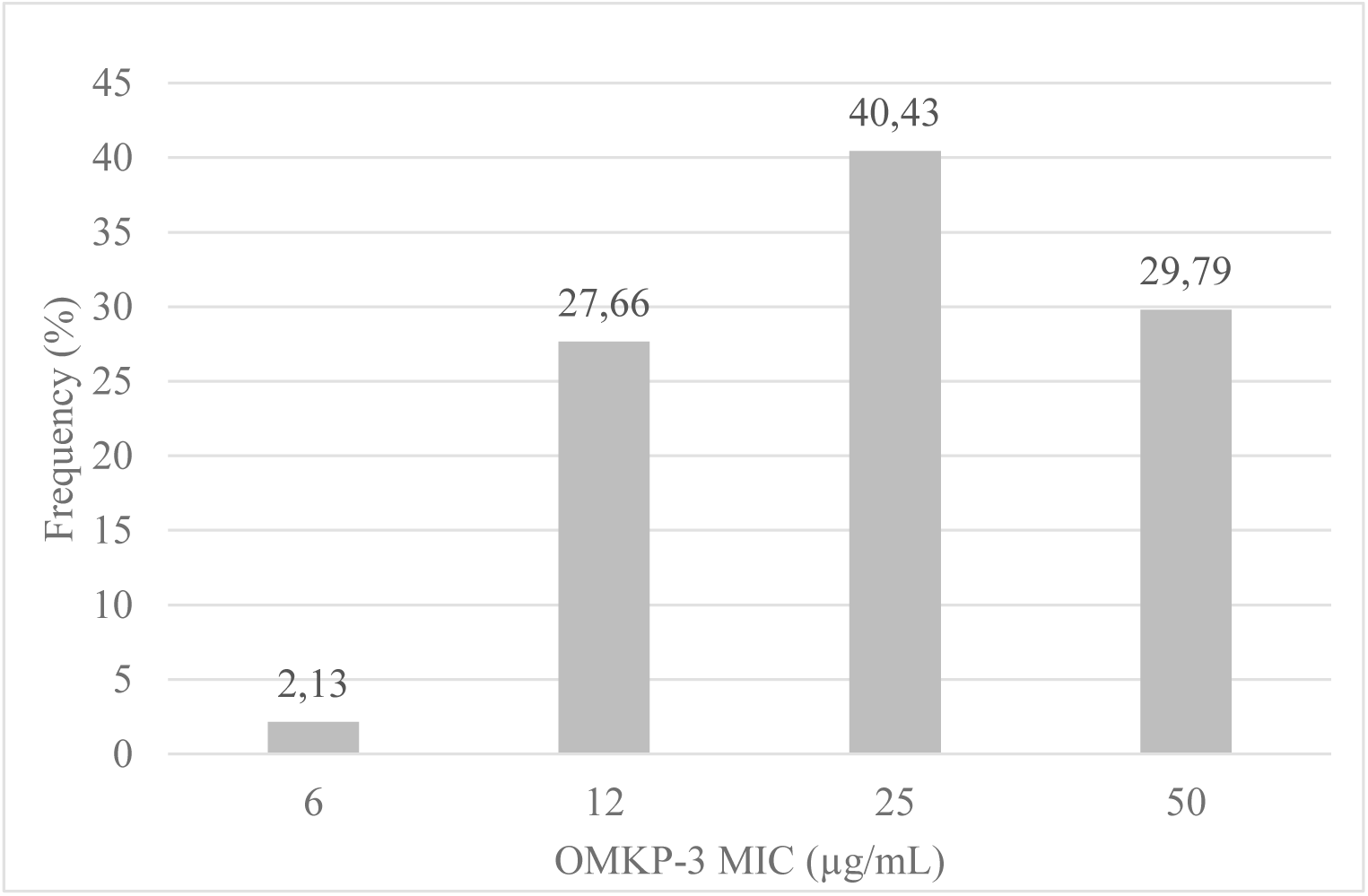
Distribution (frequency (%)) of OMKP-3 minimum inhibitory concentration (MIC) for a collection of 46 Multidrug resistant (MDR) *Klebsiella pneumoniae* clinical isolates.

**Table 3.** OMKP-3 minimum inhibitory concentrations (MIC) determination against other clinical Multidrug-resistant (MDR) Gram-negative bacterial species (*Enterobacter cloacae, Escherichia coli, Serratia marcescens and Pseudomonas aeruginosa*) in M9 minimal medium.

| Bacterial species | MIC ($\mu\text{g/mL}$) |
| --- | --- |
|  | OMKP-3 |
| *Enterobacter cloacae* (OND303) | 25 |
| *Enterobacter cloacae* (OND105e1) | 25 |
| *Escherichia coli* (OND255B) | 6.25 |
| *Escherichia coli* (OND396) | 6.25 |
| *Serratia marcescens* (OND401B) | 6.25 |
| *Pseudomonas aeruginosa* (OND162e3) | 12.5 |

### OMKP-3 exhibited antibiofilm activity against clinical MDR *K. pneumoniae*

To assess the antibiofilm capacity of OMKP-3, we evaluated OMKP-3 ability to prevent biofilm formation or inhibit pre-formed mature biofilms in MDR *K. pneumoniae* (S10p9A, OND234). OMKP-3 showed its greatest ability to prevent biofilm formation at 200µg/mL, reducing biofilm formation by 20.69% in the environmental strain S10p9A, and by 75.6% in the clinical strain OND234 (**Figure S2**). At the same concentration, the complex showed activity against already pre-formed biofilms: OMKP-3 inhibited further biofilm development by 39.3% in S10p9A strain, and by 46.2% in OND234 (**Figure S2**). However, OMKP-3 did not eradicate mature biofilms.

### OMKP-3 was non-cytotoxic *in vitr*o

To assess OMKP-3 cytotoxicity on mammalian cells, cytotoxicity assays were performed against V79 cells, an hamster lung fibroblast cell line. OMKP-3 concentrations tested ranged between 0.25xMIC and 16xMIC and cell viability was assessed by using the Thiazolyl Blue Tetrazolium Bromide (MTT) method (**Figure 3**). At 24 hours, OMKP-3 exhibited non-cytotoxicity, showing mean cell viability ratios higher than 70% for all tested concentrations. At 48 hours, OMKP-3 demonstrated non-cytotoxic values up to 4xMIC, however, minimal cytotoxicity was observed at 8xMIC and 16xMIC (cell viability of ≈60%). These results indicated that OMKP-3 was generally non-cytotoxic against V79 cells, demonstrating minimal cytotoxicity levels only at high concentrations after an exposure period of 48 hours. Noteworthy, the concentration at which OMKP-3 was active remained within the non-cytotoxic range across all tested clinical MDR *K. pneumoniae* isolates and other Gram-negative pathogens, and for the observed biofilm prevention effect.

**Figure 3.**
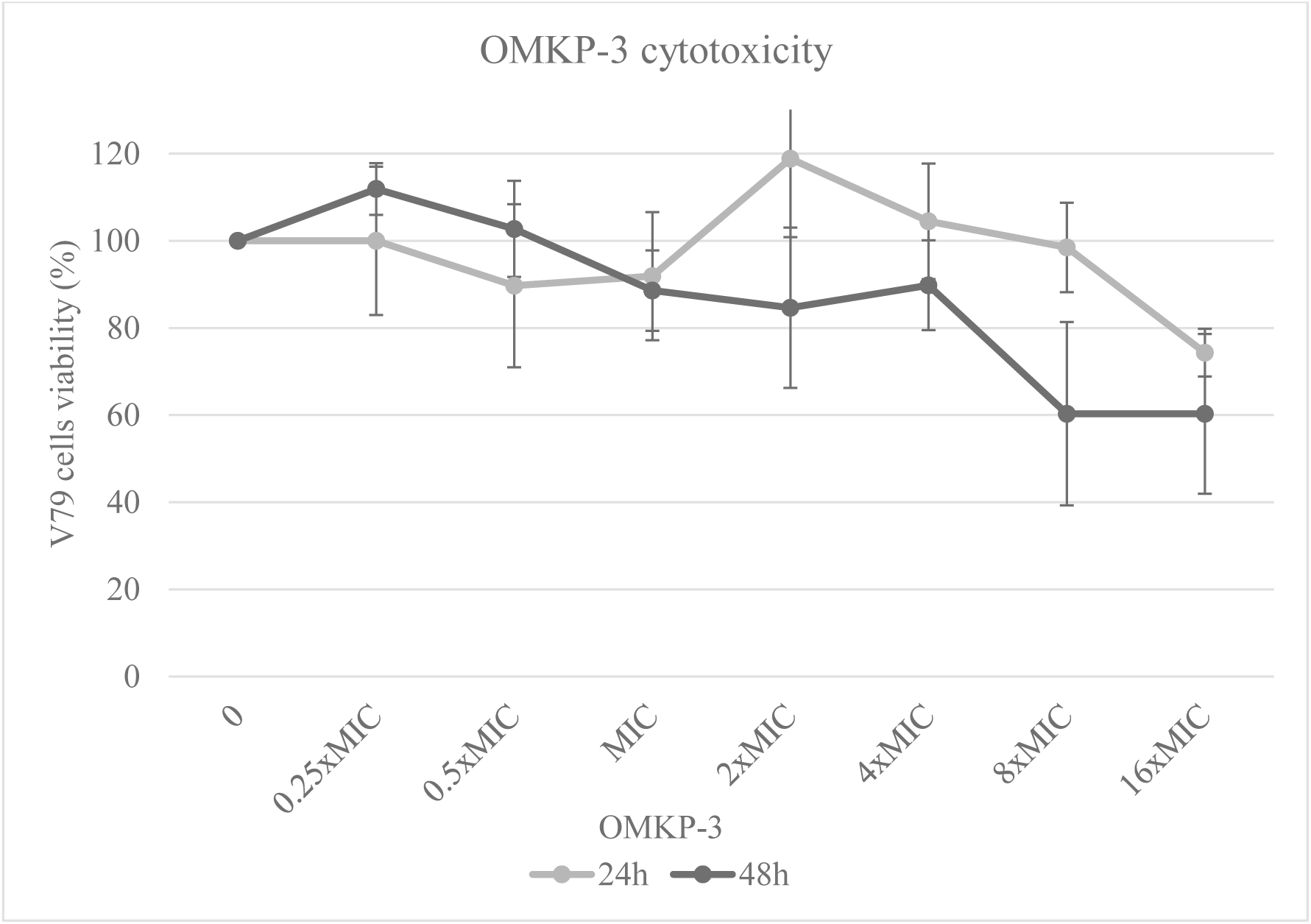
Viability of V79 cells after 24h and 48h exposure to OMKP-3 (0-16xMIC) using the MTT assays.

### OMKP-3 activity was enhanced by polymyxin B

Synergistic interactions were observed between OMKP-3 and polymyxin B (PMB) for *K. pneumoniae* S10p9A and *E. coli* ATCC25922, exhibiting a mean fractional inhibitory concentration (FIC) index of 0.38 and 0.35, respectively (**Figure 4**). This result indicates that PMB’s ability to disrupt bacterial membranes facilitated intracellular penetration of OMKP-3 inside the cell resulting in increased antimicrobial activity. Indifferent interaction between the two compounds was observed instead for OND234 *K. pneumoniae* strain, presenting a mean FIC index of 0.63 (**Figure 4**), indicating that OMKP-3 showed a strain-dependent interaction with PMB, probably due to specific bacterial characteristics of the strain.

**Figure 4.**
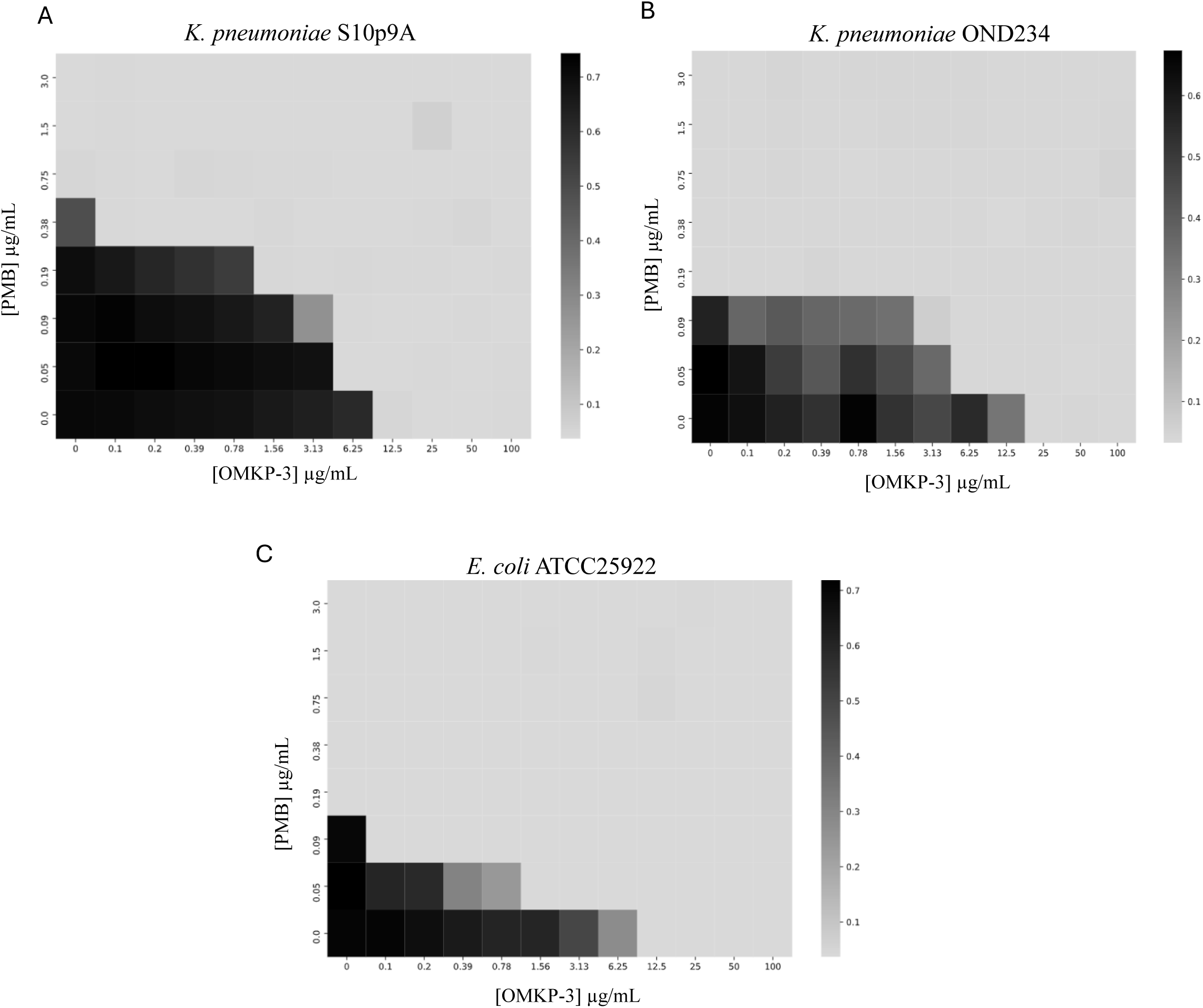
OMKP-3 (0-100µg/mL) interaction with the membrane disruptor polymyxin B (PMB) (0-3µg/mL) for *Klebsiella pneumoniae* S10p9A (A), *K. pneumoniae* OND234 (B) and *Escherichia coli* ATCC25922 (C) by checkerboard assay.

The synergistic interactions observed for the later strains further suggests that, not only OMKP-3 activity was enhanced by the action of PMB, but also OMKP-3 had the potential to act as a polymyxin B adjuvant, improving its antimicrobial activity.

### OMKP-3 mechanistic insights

#### OMKP-3 uptake may be porin-mediated

By combining the information on the distribution of MICs in the *K. pneumoniae* population (n=46) (**Table S1**) and the genomic content of the tested isolates, we found a correlation between the highest OMKP-3 MICs and the occurrence of Ompk35 and Ompk36 porin gene truncations. Our results showed that ≈43% of the isolates with an MIC of 50µg/mL exhibited *ompk35*/*ompk36* truncations, while only ≈15% of isolates with an MIC of 12µg/mL had these genes truncated (**Figure S3**). The differences between the group prevalences were not statistically significant (Fisher exact test; p-value = 0.192), however, this can derive from the fact that the sample size was small. Nevertheless, the results suggest that bacterial porins may play an important role in OMKP-3 entry across *K. pneumoniae* outer membrane, which goes accordingly with OMKP-3 chemical features.

#### OMKP-3 accumulated intracellularly without significant efflux pumps extrusion

OMKP-3 accumulation inside *K. pneumoniae* OND234 was observed by the quantification of iridium inside bacterial cells by spectrometry (ICP-AES), after OMKP-3 treatment (8xMIC, 20 minutes). Noteworthy, after treatment, OD_600nm_ was reduced by over half compared to the negative control (OD_600nm_ reduction = 56.7%), indicating that a great proportion of bacterial cells were likely already disrupted, contributing to no iridium accumulation. Iridium relative concentrations were OD_600nm_-normalized and a mean of 1.57µg/mL was determined as the OMKP-3 intracellular concentration. To prove that, despite its intracellular accumulation, OMKP-3 was not prone to be extruded by efflux pumps, we assessed the contribution of efflux pumps in OMKP-3 activity. For that we used PAβN, a commonly used efflux pump inhibitor that targets the major resistance-nodulation-division (RND) family of efflux pumps (11). PAβN was used at a subinhibitory concentration (50µg/mL), as confirmed to exert minimal antimicrobial effect (21). We observed that the presence of PAβN, had minimal to no effect on OMKP-3 activity, remaining either unchanged or two-fold lower, indicating limited efflux involvement in OMKP-3 activity.

#### OMKP-3 had a low dose range for resistance suppression

To assess the propensity of OMKP-3 to induce resistance in MDR *K. pneumoniae* we performed *in vitro* resistance development experiments using sequential passages of *K. pneumoniae* OND234 in increasing OMKP-3 concentrations (from 0.25xMIC to 16xMIC). This approach enabled the selection of spontaneous mutations associated with bacterial adaptation to OMKP-3, thereby providing insights into the potential molecular target and the resistance mechanism. To exclude the emergence of mutations arising from adaptation to growth in minimal media, parallel passaging was performed in M9 minimal media alone. Ciprofloxacin was included as a positive control, since *K. pneumoniae* OND234 was susceptible to this antibiotic, as confirmed by disk diffusion (inhibition zone diameter = 27mm).

Using this method, we generated OMKP-3-resistant mutants with a 4-fold increase in MIC. Notably, this increase was substantially lower than that obtained for ciprofloxacin (>16-fold increase), suggesting that OMKP-3 had a lower propensity to drive high-level resistance development. Additionally, resistant strains were suppressed with low antibiotic concentrations (4xMIC), which are still within the non-cytotoxic range.

#### Evidence for bacterial membranes damage caused by OMKP-3

To investigate possible targets and/or the mechanism of resistance, the whole genome of both the *K. pneumoniae* OND234 OMKP-3-resistant strain and the strain adapted to M9 minimal media (background mutational control) were sequenced and compared using the Snippy software. Whole genome sequencing showed a single missense mutation uniquely associated with the resistant strain (single-nucleotide polymorphisms - Ala54Gly) in the outer membrane stress-sensor protease/chaperon DegS. This protein contains a transmembrane domain anchored in the inner membrane and a periplasmic domain involved in sensing unfolded outer membrane proteins, thereby triggering a signaling pathway that contributes to reducing protein folding-stress issues in bacteria (22). Considering the role of this protein in the outer membrane stress response, the identified mutation is likely linked to bacterial adaptation to OMKP-3 exposure. One possible interpretation is that OMKP-3 perturbs outer membrane proteins thereby inducing the DegS sensing pathway. The Ala54Gly substitution could have modified the structure of DegS potentially increasing its protease and/or chaperone activity and enabling the mutant strain to better tolerate OMKP-3 action (**Figure 5**).

**Figure 5.**
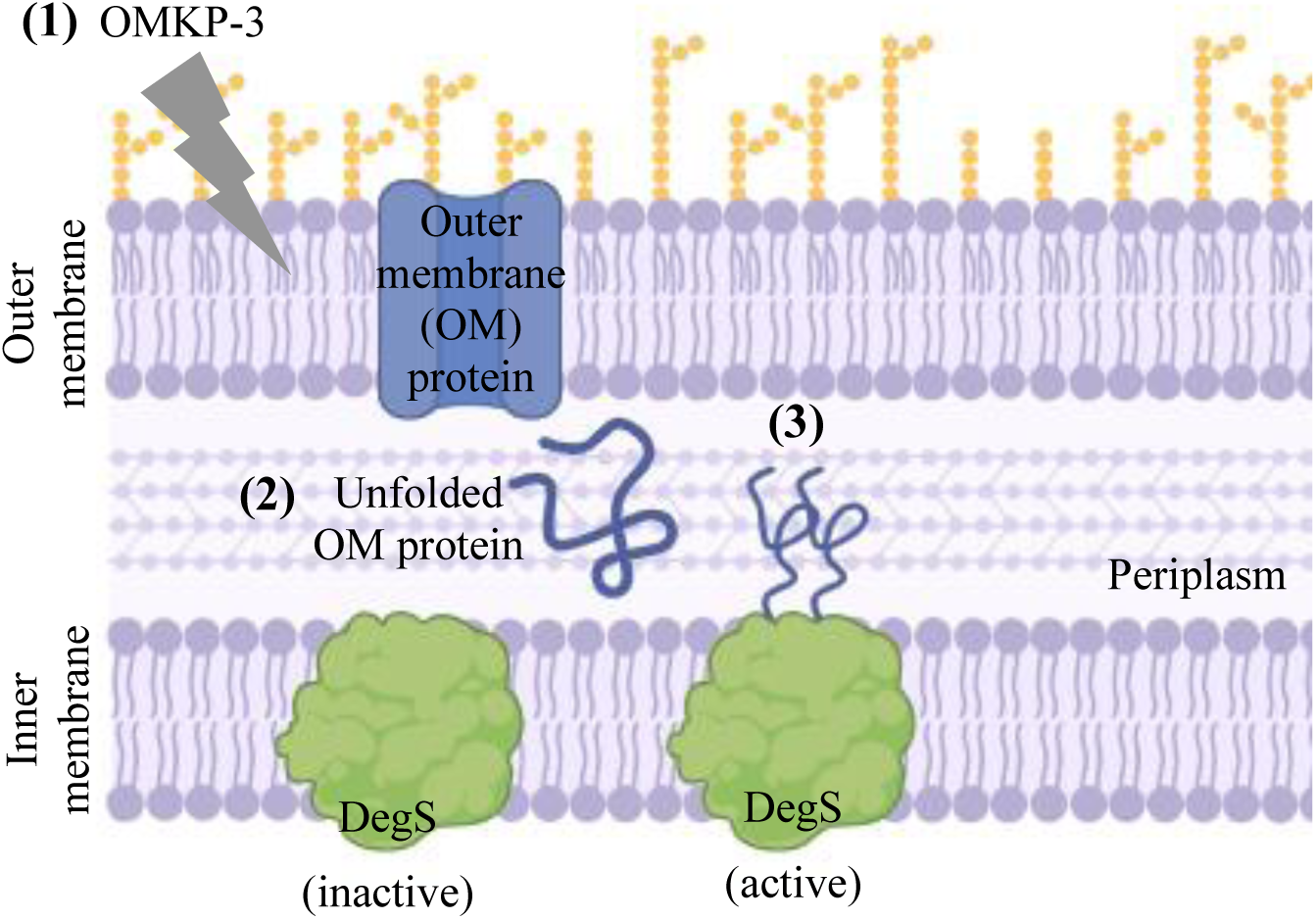
Hypothetical model of OMKP-3-mediated membrane stress in MDR *Klebsiella pneumoniae* membranes. (1) OMKP-3 may interact with the bacterial outer membrane, (2) leading to the accumulation of misfolded or unfolded outer membrane proteins (OM). (3) The presence of these defected proteins active the outer membrane stress-sensor protease/chaperone DegS which initiates a downstream cellular pathway to reduce folding-stress issues in *K. pneumoniae*. Mutations in DegS may allow *K. pneumoniae* to resist the action of OMKP-3 at low concentrations, restoring more efficiently membrane homeostasis.

To further investigate OMKP-3 possible effect on *K. pneumoniae* cell membranes we observed their integrity by fluorescence microscopy. After OMKP-3 treatment (8xMIC, 20 minutes), *K. pneumoniae* OND234 was exposed to the Live/Dead fluorophores SYTO9 and propidium iodide (PI), both nucleic acid stains (**Figure S4**). These fluorophores are commonly used to distinguish between live and dead bacteria based on the bacterial membrane integrity state, where SYTO9 stains in green and enters both viable and dead bacterial cells and PI only in membrane-permeated dead cells (23). Surprisingly, no significant SYTO9 staining was observed in the negative control (intact cells), despite the high bacterial load, indicating a natural lack of SYTO9 permeation in this *K. pneumoniae* strain. In OMKP-3 treated cells, green staining was prominently observed, indicating that OMKP-3 increased SYTO9 permeability compared with the negative control. Yet, minimal PI staining was observed. *K. pneumoniae* S10p9A strain was subjected to the same experiment to confirm the previously observed effects of OMKP-3, using a lower concentration treatment (MIC, 20min) (**Figure 6A**). Contrarily to the later strain, *K. pneumoniae* S10p9A showed greater intrinsic permeability to SYTO9, with intact bacterial cells being green stained in the negative control. Similarly, the green mean fluorescence signal was significantly higher in OMKP-3-treated cells (36.1 A.U.) compared to the negative control (14.98 A.U.) (p-value=0.002) (**Figure 6B**), further confirming that OMKP-3 have increased SYTO9 permeability across cell envelop.

**Figure 6.**
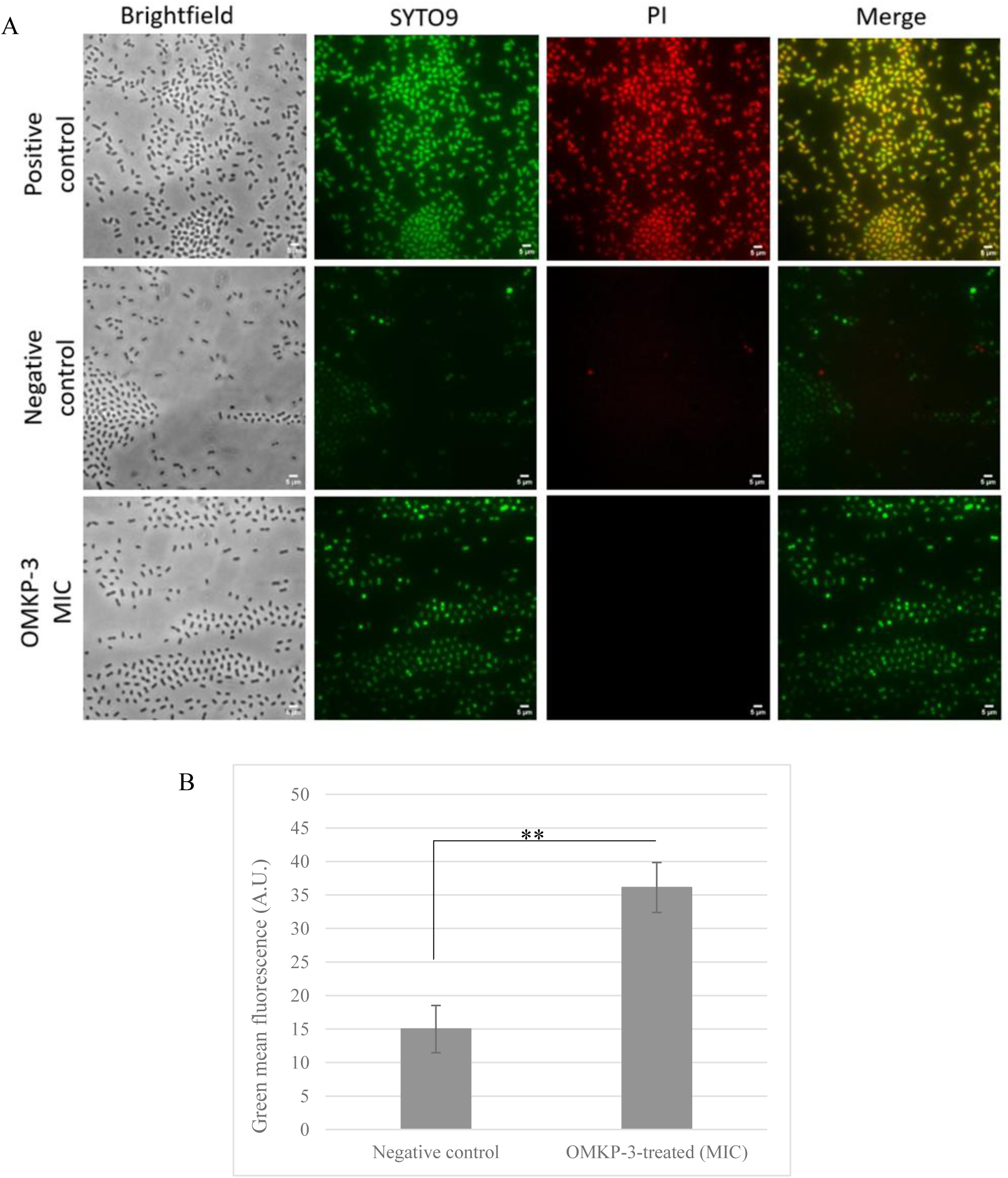
(A) Live/Dead bacterial staining images acquired from fluorescence microscopy for *Klebsiella pneumoniae* S10p9A strain after OMKP-3 treatment (MIC, 20 minutes). Positive control: boiled bacterial cells (95°C); negative control: cells treated with M9 minimal media + 0.2% DMSO. (B) Mean green (SYTO9) fluorescence measurements (Arbitrary Units (A.U.)) for *K. pneumoniae* S10p9A comparing the negative control and OMKP-3 treated cells. **p-value≤0.01 (student’s t-test).

To confirm the novelty of the OMKP-3 mode of action, we further performed cross-resistance assays by testing possible alteration in susceptibility to other common antibiotics with distinct modes of actions in the OMKP-3-resistant mutant. No phenotypic changes were observed regarding the resistance profiles of the OMKP-3 resistant mutants, when compared to the wild-type strain (**Table S2**). This result indicated that OMKP-3 resistance is not likely associated with resistance to other antibiotics through shared mechanisms, further emphasizing a possible novel membrane-related mode of action.

## Discussion

The field of antimicrobial research has been searching for new strategies to overcome the challenging rise of multidrug-resistant (MDR) infections and the traditional approaches seem to be struggling to supply this medical need for new antibiotics (14). In this worrisome scenario, the antimicrobial potential of metal-based complexes has gained significant attention in the past few years, since they represent a novel strategy for the development of antimicrobial agents (15).

Here we studied the antimicrobial activity of a novel iridium-based metal complex, OMKP-3. We found that OMKP-3 exhibited robust antimicrobial activity against carbapenem-resistant and ESBL-producing *K. pneumoniae* strains, of clinical and food-chain origins, and susceptible *E. coli* reference strain in M9 minimal media. When tested in rich growth media (such as MHB), OMKP-3 lost its antimicrobial activity, a phenomenon previously described for other metal-based antimicrobial complexes (24), highlighting the importance of evaluating metal-based antimicrobials under different nutrient conditions. In this study we further confirmed that the loss of activity was directly associated with the presence of a high load of amino acids in the media, as already hypothesized to be a contributing factor (19). Importantly, the antimicrobial activity of OMKP-3 remained unchanged in the presence of low amino acid concentrations compared to rich bacterial media. Although these conditions more closely approximate physiological environments (25), they remain sub-physiological, and thus warrant further evaluation under more representative *in vivo*–like conditions.

Furthermore, M9 minimal medium was previously described to better mimic the *in vivo* lung conditions, where amino acid levels are minimal, reflecting more realistically the performance of antimicrobials in infection scenarios (26). Moreover, we confirmed that OMKP-3 potent antimicrobial activity in M9 minimal media was a characteristic of the complex, and not a broad response to nutritional stress by bacteria, as shown by the unchanged or even reduced antimicrobial activity of other commonly used antibiotics in this minimal media. OMKP-3 exhibited potent bactericidal activity against MDR *K. pneumoniae* and susceptible *E. coli*, achieving an even lower MBC/MIC ratio than that for cefiderocol (27), a recently approved antibiotic with broad activity against carbapenem-resistant Gram-negative bacteria (28). OMKP-3 was able to completely eradicate bacterial cultures at 8xMIC within a maximum of two hours, where over half of the bacterial cell density was reduced after only 20 minutes of exposure. Additionally, OMKP-3 killing curves enabled the identification of a concentration-dependent kinetic. The rapid bactericidal activity of OMKP-3 further suggests disruption of essential bacterial functions or components needed to maintain cellular integrity, such as DNA or membrane damage (29), or the production of lethal doses of hydroxyl radicals, killing bacteria by oxidative stress (30), an hypothesis that will be further explored for OMKP-3.

OMKP-3 spectrum of activity was assessed against 46 clinical carbapenem-resistant *K. pneumoniae* isolates recovered from invasive infections and with varied genetic backgrounds (20). OMKP-3 inhibited all tested clinical isolates at non-cytotoxic concentrations, confirming this complex as a promising antimicrobial against MDR *K. pneumoniae*. However, activity was achieved at moderately high drug concentration for some isolates in the population, indicating that further optimization may be warranted. In the studied bacterial population, we observed that higher OMKP-3 MICs were generally associated with the existence of Ompk35 and Ompk36 porins gene truncations. These results suggest that porins may be an important mean of OMKP-3 entry in *K. pneumoniae* and that MIC variations may derive from porin-mediated variation in membrane permeability. This finding goes accordingly with OMKP-3 chemical properties, as small polar molecules (≈600-700 Da) are usually able to cross through bacterial porins (11, 31). Importantly, no association was observed between higher MICs and the carriage of antimicrobial resistance determinants, virulence genes or epidemiological background in the MDR *K*. *pneumoniae* isolates analysed. Nevertheless, we cannot discard that other unidentified strain-specific characteristics may be involved in the variation of OMKP3-activity in the population.

We demonstrated that OMKP-3 antimicrobial activity was not restricted to *K. pneumoniae*, as it was active against other Gram-negative MDR human pathogens, including *Enterobacter cloacae, E. coli, Serratia marcescens and Pseudomonas aeruginosa*, suggesting a broad-spectrum activity and a potentially shared mechanism of action and/or molecular targets among Gram-negative bacteria. OMKP-3 demonstrated antibiofilm activity against MDR *K. pneumoniae* having prevented formation and inhibited growth within mature biofilm. This feature should be further investigated for its application as prophylactic or anti-adhesion agent, especially considering its high activity against the clinical strain tested.

OMKP-3 was non-cytotoxic against mammalian cells (V79 cell line), exhibiting no significant cytotoxicity levels after 24 hours of incubation, even at concentrations as high as 16xMIC. Nonetheless, further studies in infection models are needed to prove its efficacy *in vivo*.

The synergistic interaction observed between OMKP-3 and polymyxin B suggests that the capacity of polymyxin to disrupt outer and inner membrane (32) may facilitate OMKP-3 penetration across cell envelop and consequently enhancing its antimicrobial activity. This interaction indicates that OMKP-3 penetration inside bacterial cell may be a crucial step for its antimicrobial activity. Importantly, OMKP-3 improved the antimicrobial activity of polymyxin B acting as an antibiotic adjuvant, a clinically significant result. Little can be inferred regarding OMKP-3 mechanism of action by the synergistic interactions with polymyxin B since many antibiotics with different targets can commonly synergise with polymyxin B (33). The synergistic-indifferent interactions differences between the two *K. pneumoniae* strains tested further suggests different membrane compositions, or even differences in permeability and efflux among the tested strains (34), therefore, this interaction should be further investigated for a higher number of *K. pneumoniae* strains. Furthermore, evaluating its synergistic activity in combination with other antibiotic classes is warranted.

We proved that OMKP-3 was able to accumulate intracellularly in the carbapenem-resistant *K. pneumoniae* strain tested, therefore, overcoming the challenging and highly impermeable outer membrane in these bacteria. OMKP-3 uptake was fast and effective, as even reduced intracellular concentrations were sufficient to exert a potent antimicrobial activity leading to a large reduction in bacterial biomass. Despite its intracellular accumulation, OMKP-3 exhibited minimal involvement of RND efflux pumps, as indicated by checkerboard assays with RND efflux pump inhibitors, suggesting that its chemical structure is poorly recognized as a substrate by these transporters. OMKP-3 contains a primary amine, is amphiphilic and has a positive charge in solution, chemical features associated with improved accumulation and less susceptibility to efflux in Gram-negative bacteria, according to the proposed eNTRy rules (35, 36). Yet, other chemical properties such as a low tridimensionality and high rigidity (35), characteristics that haven’t been fully characterized for OMKP-3, may further contribute to the observed intracellular accumulation and minimal efflux involvement. Importantly, when compared with other commonly used antibiotics including β-lactams, chloramphenicol and erythromycin tested against susceptible *K. pneumoniae* ATCC11296 and MDR strains (minimal 4-fold to 16-fold MIC decrease in the presence of efflux inhibitor), OMKP-3 showed lower involvement of efflux mechanisms (37).

OMKP-3 resistance propensity was compared with ciprofloxacin by sequential passages in increasing concentrations of both antimicrobials. Compared to the high resistance level obtained for ciprofloxacin (>16-fold MIC increase), the low fold change induced by OMKP-3 (4-fold MIC) suggests that low and still non-cytotoxic concentrations can be used to kill resistant bacteria. Despite the promising results, further experiments need to be performed to confirm the low propensity of OMKP-3 for antimicrobial resistance. Importantly, no cross-resistance was observed between OMKP-3 and other commonly used antibiotics from different classes, an indication that OMKP-3 resistance mechanism/target, is not shared with other antibiotics. In the OMKP-3-resistant strain, one single missense mutation was found in an outer membrane stress-sensor protease/chaperone DegS, likely involved in the acquired low-level OMKP-3 resistance. This mutation suggests an involvement of OMKP-3 with *K. pneumoniae* membranes, as DegS is activated by unfolded outer membrane proteins (22). OMKP-3 effect on bacterial membranes was further evidenced by fluorescent microscopy using the Live/Dead fluorophores SYTO9 and propidium iodide (PI). Contrarily to the untreated bacterial cells that showed limited SYTO9 fluorescence signal, exposure to OMKP-3 (MIC, 8xMIC) allowed for an increased SYTO9 signal in both MDR *K. pneumoniae* strains. These results indicate a higher SYTO9 permeability across cell envelop and further evidence OMKP-3 involvement with bacterial membranes. This observation goes accordingly with previous reports on Gram-negative bacteria, where a stronger green fluorescence signal was observed for dead *P. aeruginosa* bacterial cells treated with isopropanol compared with untreated cells (23). However, in both strains, minimal PI staining was observed, even with a high concentration treatment where bacterial density was reduced by half compared to the negative control. These findings suggest that OMKP-3-induced membrane alterations allowed for enhanced SYTO9 permeability, yet, these were insufficient for the larger molecule PI to penetrate and confer the characteristic ‘dead’ phenotype obtained by this Live/Dead kit. Alternatively, OMKP-3-induced killing and bacterial lysis may occur so rapidly that PI does not have sufficient time to enter cells and bind DNA before cell disruption.

Altogether, our results, point towards a membrane-related mode of action and resistance mechanism, yet the involvement of other OMKP-3 molecular targets should be further explored. Moreover, it remains to be clarified if the putative resistance mechanism suggested here coincides with the drug target and further genetic studies are needed to prove the causality between the *degS* mutation and the resistance phenotype.

## Material and Methods

### Ethical statement

Bacterial isolates (OND) were obtained in a surveillance study done in five hospitals in the region of Lisbon, under the scope of project ONEIDA (20) that was approved by the hospital’s ethical committees (38). *Klebsiella pneumoniae* S10p9A strain here tested was isolated from a sample collected as part of the routine processing control at a slaughterhouse, under the supervision of a veterinarian and with the oral consent of the slaughterhouse management. The Ethical Committee approved the study for Research and Teaching of the Faculty of Veterinary Medicine, Universidade de Lisboa (N/Refª 038/2020).

### OMKP-3 chemical synthesis and storage

OMKP-3 was synthesized under an inert nitrogen atmosphere using standard Schlenk techniques, following a procedure previously reported by some of the authors (18). Briefly, the appropriate imidazolium salt, NHC proligand (0.125 g, 0.3 mmol) was combined with silver oxide (Ag_2_O; 0.051 g, 0.225 mmol) in dichloromethane (CH_2_Cl_2_, 5 mL) and stirred at room temperature for 2h. Subsequently, the metal precursor pentamethylcyclopentadienyl iridium dichloride dimer (0.12 g, 0.15 mmol) was added, and the reaction mixture was stirred at room temperature overnight. The resulting suspension was filtered through Celite, and the solvent was removed under reduced pressure. The crude product was purified by silica gel column chromatography using a dichloromethane/acetone mixture.

Structural characterization was performed by ^1^H and ^13^C nuclear magnetic resonance (NMR) spectroscopy on a Bruker Avance III 400 MHz spectrometer. The ^1^H and ^13^C NMR spectra were in agreement with the data previously reported for OMKP-3 (18). High-resolution mass spectrometry (HRMS) was conducted in positive ion mode using Thermo Scientific Focus hybrid quadrupole-Orbitrap mass spectrometer, and the obtained mass spectrum was likewise consistent with previously reported values. OMKP-3 was stored in powdered form protected from light at 4°C. Stock solutions (50mg/mL) were prepared in 100% dimethyl sulfoxide (DMSO), and stored at -20°C, protected from light. The stock concentration was selected to ensure that the final DMSO concentration in biological assays did not exceed 2.5%, thereby avoiding DMSO-associated toxicity.

### Bacterial clinical isolates and control strains

Multidrug-resistant (MDR) *Klebsiella pneumoniae* isolates used in this study were previously characterized (38). Most of the analysis were performed with: *K. pneumoniae* strain S10p9A, isolated in 2019 from the pork processing chain in the Lisbon region, which is an ESBL producing strain, that belongs to sequence type (ST) 60; and OND234 strain collected in 2018 from invasive infection in humans in a Lisbon region hospital, which is a carbapenem-resistant isolate carrying KPC-3 that belongs to ST13. Additional 46 clinical isolates of MDR *K. pneumoniae* from multiple STs (n=23) (**Table S1**) were also included for determining the concentration at which 50% (MIC50) and 90% (MIC90) of the isolates were inhibited (see below). *Escherichia coli* strain ATCC25922 was used as a control for antimicrobial susceptibility testing assays. Furthermore, *Enterobacter cloacae* (OND303, OND105e1), *E. coli* (OND255B, OND396), *Serratia marcescens* (OND401B) and *Pseudomonas aeruginosa* (OND162e3) collected under the scope of project ONEIDA (38) were included to assess OMKP-3 spectrum of action.

### Determination of minimum inhibitory concentrations (MIC) and minimum bactericidal concentrations (MBC)

OMKP-3 MIC was determined by broth microdilution assays according to EUCAST guidelines (39). Briefly bacterial cultures (5x10^5^ CFU/mL) were exposed to OMKP-3 (0.19-200µg/mL) and incubated at 35°C in static aerobic conditions for 18±2 hours. After incubation, the MIC of all antimicrobials tested was recorded as the lowest concentration that inhibited visible bacterial growth (39). Three technical replicas were performed for each strain tested. To determine bacterial viability, bacterial cultures were further serially diluted and plated in MHA II (Becton Dickinson, Sparks, MD, USA) for further colony counting. The MBC was recorded as the lowest concentration causing a ≥3 log10 reduction in CFU/mL, of the viable bacteria after incubation for 18±2 hours (40). MIC50 and MIC90 were determined through MIC characterization of a diverse collection of 46 clinical MDR *K. pneumoniae* (**Table S1**).

### OMKP-3 kinetic of killing

The time-kill assay was performed according to the Clinical and Laboratory Standards Institute (CLSI) recommendations (40). Briefly, a bacterial suspension (5x10^5^ CFU/mL) was prepared in M9 minimal media (M9 salts 1x, glucose 0.4%, MgSO4 1 mM, CaCl2 0.1 mM) and exposed to OMKP-3 solutions with concentrations ranging from MIC to 8xMIC. Bacterial cultures were incubated at 37°C under constant shaking (≈200 rpm) for 24h. At the selected timepoints (0, 2, 4, 6, 8, 24h) bacterial cultures were plated on MHA II and incubated at 35°C overnight for colony counting.

### Determination of antibiofilm activity of OMKP-3

The OMKP-3 ability to inhibit or eradicate biofilms was tested according to previously described antibiofilm assays (41–43) by growing a 1:100 dilution of an overnight bacterial culture in M9 minimal media and incubating for 16-18h at 37°C without shaking. Planktonic cultures were removed and OMKP-3 solutions (0.78-200µg/mL) were dispensed on the top of the pre-formed biofilms. The cultures were incubated for 16-18h at 37°C without shaking, and the biofilm was quantified by the crystal violet staining method (44). Two biological and three technical replicas were performed for each strain tested. For the ability of OMKP-3 to prevent biofilm formation, a 1:50 dilution from an overnight culture was incubated for 16-18h at 37°C without shaking in the presence of different OMKP-3 concentrations (0.19-200µg/mL) and the biofilm was quantified by the crystal violet staining method (44). Two biological and three technical replicas were prepared.

### OMKP-3 cross-inhibition assays with an efflux inhibitor and a membrane disruptor

Interactions between OMKP-3 and membrane disruptor (Polymyxin B (PMB)) (Apollo Scientific, Stockport, UK) were verified by checkerboard synergy assays (45). Solutions of PMB (0.05-3µg/mL) and OMKP-3 (0.1-100µg/mL) were prepared in M9 minimal media and combined in a 96-well microtiter plate in a checkerboard pattern and the bacterial suspension was added (final concentration of 5x10^5^ CFU/mL). The cultures were incubated at 35°C for 18±2 hours. Drug-drug interactions were assessed by calculating the fractional inhibitory concentration (FIC) index, where the interaction was considered: synergistic if FIC≤0.5, indifferent if FIC>0.5 - <2 and antagonistic if FIC≥4 (45). Interactions between OMKP-3 and the efflux pumps inhibitors (phenylalanine-arginine beta-naphthylamide (PAβN)) (MP Biomedicals, Solon, OH, USA) were verified using the same methodology, at a fixed PaβN concentration of 50µg/mL. Three biological replicas were performed for each interaction

### OMKP-3 cytotoxicity assays

OMKP-3 cytotoxicity was assessed against the V79 cell line (Chinese hamster lung fibroblasts) and using the Thiazolyl Blue Tetrazolium Bromide (MTT) method (46). V79 cells were grown as monolayers in Dulbecco’s modification of Eagle’s medium (DMEM), supplemented with 10% FBS (fetal bovine serum), in a humidified incubator with 5% CO_2_ saturation at 37 °C. A cell suspension (3 x 10^4^ cells/mL) was seeded for 48 hours in a microtiter plate and seeded cells were exposed to OMKP-3 concentrations ranging from 200µg/mL to 3.12µg/mL, for 24 and 48 hours. MTT was added to a final concentration of 0.5mg/mL and incubated for 2 hours at 37°C. The formed precipitate was solubilized using DMSO for 15 minutes in the dark and quantified at an OD_570nm_. V79 cell viability was obtained by the quantification of MTT in OMKP-3 treated cells compared to the negative control (untreated cells in DMEM supplemented with 0.5% DMSO) and expressed in percentage (%). The complex was considered non-toxic when cell viability was ≥70% (47).

### Assessment of membrane permeability by fluorescence microscopy

A bacterial culture at OD_600nm_ of 0.8 was treated with OMKP-3 (MIC, 8xMIC) for 20 minutes and incubated at 37°C under constant shaking. The bacterial suspension was centrifuged (13000 rpm,1 min) (Heraeus Biofuge pico, Thermo Electron Corporation, Langenselbold, Germany) and cells were washed twice with NaCl 0.8% (m/v) and incubated with propidium iodide (30µM) (Biotium, Fremont, CA, USA) and SYTO9 (5mM) (Thermo Fisher Scientific, Wilmington, DE, USA) fluorophores (10 min in the dark). Bacteria were centrifuged (13000 rpm,1 min) and resuspended in 50µL of the supernatant. A volume of the bacterial suspension was dispensed over a thin layer of solidified agarose (1.7%) (LE Agarose, SeaKem, Rockland, ME, USA). In parallel, a positive control (bacterial cells boiled at 95° C for 10 min) and a negative control sample (bacterial cells treated with M9 minimal media + 0.2% DMSO) were prepared. The samples were observed by fluorescence microscopy on Leica DM 6000B upright microscope. Three technical replicas and two biological replicas were prepared. Mean fluorescence values were determined for *K. pneumoniae* S10p9A strain using ImageJ software 1.53c.

### Quantification of Iridium uptake by bacterial cells by inductively coupled plasma-atomic emission spectroscopy (ICP-AES)

Whole-cell lysates were obtained by acidic lysis as described (48). Briefly, a culture at OD_600nm_ of 0.8 was exposed to OMKP-3 at 8xMIC for 20 minutes. The bacterial suspension was centrifuged at 8000 rpm (Heraeus Biofuge pico, Thermo Electron Corporation, Langenselbold, Germany) for 10 minutes, the cells were washed once with Ethylenediaminetetraacetic acid (EDTA) (10mM) and twice with PBS1x and the OD_600nm_ recorded. Cells were centrifuged (8000 rpm, 10 minutes) and digested with 65% HNO_3_ at room temperature (25°C) for 24 hours. After digestion, PBS 1x was added, samples were centrifuged at 3600 rpm for 5 minutes and the supernatant was stored at -20°C. All samples were analysed by ICP-AES (Horiba Jobin Yvon inductively coupled plasma atomic emission spectrometer) (Analytical Laboratory, Department of Chemistry, FCT-NOVA) for iridium detection. Three technical and two biological replicas were performed. A negative control was prepared in parallel wherein OMKP-3 was substituted by M9 minimal media.

### Induction of OMKP-3 resistance

To evaluate the ability of *K. pneumoniae* to develop resistance to OMKP-3, *K. pneumoniae* OND234 was serially grown in M9 minimal media with increasing OMKP-3 concentrations ranging from 0.25xMIC to 16xMIC overnight at 37°C with constant shaking (170 rpm) (New Brunswick Innova 4300 Incubator Shaker, Eppendorf, Hamburg, Germany). In parallel, the same passages were performed in M9 minimal media (background mutational control), and in increasing ciprofloxacin concentrations (Bayer, St. Kankakee, IL, USA) (0.25xMIC to 16xMIC) (positive control). Each culture was daily stored at -80°C.

### Assessment of cross resistance between OMKP-3 and other antibiotics

Antimicrobial susceptibility of OMKP-3-resistant *K. pneumoniae* OND234 strain to other antibiotics was evaluated by Kirby-Bauer disk diffusion method, according to the EUCAST guidelines (49). The following antibiotics were tested: ticarcillin (TIC, 75 µg), temocillin (TEM, 30 µg), amoxicillin/clavulanic acid (AMC, 20/10 µg), piperacillin/tazobactam (TZP, 30/6 µg), trimethoprim/sulfamethoxazole (SXT, 1.25/23.75 µg), cefoxitin (FOX, 30 µg), cefotaxime (CTX, 30 µg), ceftazidime (CAZ, 14 µg), ceftazidime/avibactam (CZA, 30 µg), imipenem (IMP, 10 µg) and gentamicin (CN, 10 µg), ciprofloxacin (CIP, 5 µg). Strain was classified as susceptible (S), susceptible to increased exposure (I) and resistant (R), according to EUCAST guidelines (50).

### OMKP-3 resistant-mutant analysis

Total genomic DNA of the *K. pneumoniae* OND234 OMKP-3-resistant mutant and M9-adapted strains was extracted using the Dneasy® Blood and Tissue kit (QIAGEN, Hilden, Germany), according to manufacture instructions. Libraries were constructed using the Nextera XT Library Preparation Kit (Illumina, San Diego, CA, USA), and pair-ended 2x150 bp reads were produced using the NextSeq 500 platform (Illumina, 100x coverage). Raw reads were assembled using INNuca pipeline v3.1 (https://github.com/B-UMMI/INNUca) (51). The contigs resulting from the assembly were annotated using Prokka version 1.13.3 (52). The genomic differences between the reference and the mutant’s genome were analysed using Snippy software version 4.6.0 (53). The genes containing the missense mutations were identified and information regarding protein’s active/binding sites were identified using UniProt (54).

## Acknowledgements

This work was funded by the Innovalley Proof-of-concept award 2^nd^ and 3^rd^ Editions. This work was supported by FCT – Fundação para a Ciência e a Tecnologia, I.P., through MOSTMICRO-ITQB R&D Unit (doi.org/10.54499/UID/04612/2025, UID/PRR/4612/2025) and LS4FUTURE Associated Laboratory (DOI 10.54499/LA/P/0087/2020). This work was partially supported by PPBI - Portuguese Platform of BioImaging (PPBI-POCI-01-0145-FEDER-022122) co-funded by national funds from OE - “Orçamento de Estado” and by european funds from FEDER - “Fundo Europeu de Desenvolvimento Regional.

Raquel L. Almeida performed most of experimental work, analyzed the data and wrote the manuscript. Oscar Lenis-Rojas designed and synthesized the studied complex, analyzed data, supervised the work and reviewed the manuscript. Mariana Araújo contributed to preliminary experimental work and reviewed the manuscript. Claudia Malta-Luís performed the cytotoxicity assays and reviewed the manuscript. Francisco C. Mendes contributed to complex synthesis and reviewed the manuscript. Beatriz Royo, provided resources and facilities, designed the metal complex, supervised the work, analyzed results and reviewed the manuscript. Maria Miragaia provided resources and facilities, conceptualized the project, designed the experimental approach, supervised the work, and reviewed the manuscript. All authors approved the final version.

## Supplementary Material

**Figure S1** – OMKP-3 chemical structure.

**Figure S2** – OMKP-3 antibiofilm ability against MDR *Klebsiella pneumoniae* strains S10p9A and OND234: (A) OMKP-3 biofilm prevention ability (%); (B) OMKP-3 mature biofilm inhibition (%).

**Figure S3** – Porin (Ompk35 and Ompk36) gene truncation prevalence (%) among MDR *Klebsiella pneumoniae* clinical isolates according to their OMKP-3 minimum inhibitory concentration (MIC) (12.5 µg/mL or 50 µg/mL).

**Figure S4** – Live/Dead bacterial staining images acquired from fluorescence microscopy for *Klebsiella pneumoniae* OND234 strain after OMKP-3 treatment (MIC, 20 minutes). Positive control: boiled bacterial cells (95°C); negative control: cells treated with M9 minimal media + 0.2% DMSO.

**Table S1** – Information previously obtained for the Multidrug-resistant *Klebsiella pneumoniae* isolates (n=46) – hospital of origin, biological collection site, sequence type (ST), resistance genotype, presence of porin (Ompk35 and Ompk36) gene truncation (expressed in percentage of amino acid length from the start codon) – in combination with the correspondent OMKP-3 minimum inhibitory concentration (MIC). Frequency (%) of Ompk35 and Ompk36 porin gene truncation in *K. pneumoniae* population. AMG: Aminoglycosides; TMP-SUL: Trimethoprim-sulfamethoxazole; FQL: Fluroquinolones; CHL: Chloramphenicol RIF; Rifampicin; TET; tetracycline; COL: colistin; MAC: Macrolides.

**Table S2** – Cross-resistance assessment between OMKP-3 and other common antibiotics from different classes (aminoglycosides, fluoroquinolones, cephalosporins, carbapenems, penicillins and miscellaneous) by disk diffusion method (halo diameter determination) for wild-type (WT) and OMKP-3-resistant mutant *K. pneumoniae* OND234. Strain was classified as susceptible (S), susceptible to increased exposure (I) and resistant (R), according to EUCAST guidelines (2024).

