## Supplementary data for "*In Vitro* Activity of a Novel Metal-Based Antimicrobial against Multidrug-Resistant *Klebsiella pneumoniae*"

**Table S1** – Information previously obtained for the Multidrug-resistant *Klebsiella pneumoniae* isolates (n=46) – hospital of origin, biological collection site, sequence type (ST), resistance genotype, presence of porin (Ompk35 and Ompk36) gene truncation (expressed in percentage of amino acid length from the start codon) – in combination with the correspondent OMKP-3 minimum inhibitory concentration (MIC). Frequency (%) of Ompk35 and Ompk36 porin gene truncation in *K. pneumoniae* population. AMG: Aminoglycosides; TMP-SUL: Trimethoprim-sulfamethoxazole; FQL: Fluroquinolones; CHL: Chloramphenicol RIF: Rifampicin; TET; tetracycline; COL: colistin; MAC: Macrolides.

| *K. pneumoniae* strain | Hospital | Biological collection site | Sequence type (ST) | Antimicrobial resistance genotype | OMKP-3 MIC (µg/mL) | Ompk35 and Ompk36 porin gene truncation | Frequency of porin gene truncation (%) |
|---|---|---|---|---|---|---|---|
| OND337 | C | urine | 29 | AMG; b-lactams | 6,25 | - | - |
| OND207 | A | urine | 15 | AMG; b-lactams | 12,5 | - | 15.4 |
| OND208 | A | urine | 348 | AMG; b-lactams; TMP-SUL | | - | |
| OND209 | A | urine | 15 | AMG; b-lactams; TMP-SUL; FLQ | | - | |
| OND215A | A | urine | 17 | AMG; b-lactams; TMP-SUL; FLQ; RIF | | - | |
| OND253 | A | pus | 13 | AMG; b-lactams; TMP-SUL; CHL | | OmpK35-70% | |
| OND258 | A | urine | 348SLV | AMG; b-lactams; TMP-SUL; FQL | | - | |
| OND270 | B | urine | 1142 | AMG; b-lactams; SUL; FQL; CHL | | - | |
| OND334 | C | rectal swab | 13 | AMG; b-lactams; TMP-SUL; FQL; CHL; TET | | - | |
| OND436 | E | pus | 348 | AMG; b-lactams; TMP-SUL; FQL; COL | | - | |
| OND439 | E | biopsy | 13SLV | AMG; b-lactams; TMP-SUL | | - | |
| OND445 | E | rectal swab | 3403SLV | AMG; b-lactams; TMP-SUL; FQL | | - | |
| OND430 | E | rectal swab | 323 | AMG; b-lactams; TMP-SUL | | - | |
| OND204 | A | urine | 147 | AMG; b-lactams; TMP-SUL; FQL; RIF | 25 | - | 31 |
| OND205 | A | haemoculture | 257TLV | AMG; b-lactams; TMP-SUL | | - | |
| OND210 | A | urine | 45 | AMG; b-lactams; TMP-SUL; FQL; TET; CHL | | - | |
| OND221A | A | catheter | 231 | AMG; b-lactams; TMP-SUL; CHL; TET; RIF | | - | |
| OND236 | A | urine | 34 | AMG; b-lactams; TMP-SUL | | - | |
| OND242A | A | bronchial secretions | 459 | AMG; b-lactams; TMP-SUL; FQL | | OmpK35-28% | |
| OND249 | A | haemoculture | 3650 | AMG; b-lactams; TMP-SUL | | - | |
| OND261 | B | blood | 323 | AMG; b-lactams; TMP-SUL; FQL; CHL; TET | | OmpK35-3% | |
| OND262 | B | urine | 147 | AMG; b-lactams; TMP-SUL; FQL; CHL; RIF; MAC | | - | |
| OND283 | B | rectal swab | 323 | b-lactams; TMP-SUL; FQL; CHL; TET; COL | | OmpK35-3% | |
| OND306 | C | rectal swab | 14 | AMG; b-lactams; TET; COL | | OmpK36-0% | |
| OND317 | C | rectal swab | 392 | AMG; b-lactams; TMP-SUL; FQL; CHL; TET | | - | |
| OND318 | C | pus | 348 | AMG; b-lactams; TMP-SUL; FQL | | OmpK36-7% | |
| OND394 | D | bronchial wash | 307 | AMG; b-lactams; TMP-SUL; FQL; CHL; TET; COL | | - | |
| OND405 | E | ascitic fluid | 2217 | AMG; b-lactams; TMP-SUL | | - | |
| OND416 | E | cephalorachidian fluid | 1229 | AMG; b-lactams; TMP-SUL; TET | | - | |
| OND431 | E | urine | 4585 | b-lactams; COL | | - | |
| OND260 | B | urine | 231 | AMG; b-lactams; TMP-SUL; CHL; RIF; TET | | - | |
| OND419 | E | rectal swab | 17 | AMG; b-lactams; TMP-SUL; FQL; CHL; RIF; MAC; COL | | OmpK36-84% | |
| OND203 | A | bronchial secretions | 35 | AMG; b-lactams; TMP-SUL; FQL; CHL; RIF | 50 | - | 42.9 |
| OND214A | A | prosthetic | 2202SLV | AMG; b-lactams; TMP-SUL | | - | |
| OND248 | A | sputum | 17 | AMG; b-lactams; TMP-SUL; FQL; CHL; RIF; MAC | | OmpK36-30% | |
| OND259 | B | blood | 17 | AMG; b-lactams; TMP-SUL; FQL; CHL; RIF; MAC | | - | |
| OND308 | C | rectal swab | 35 | AMG; b-lactams; TMP-SUL; FQL; CHL; RIF | | - | |
| OND316A | C | rectal swab | 14 | TET | | OmpK36-33% | |
| OND402 | E | rectal swab | 2944 | b-lactams; TMP-SUL; FQL | | OmpK36-70% | |
| OND404 | E | urine | 231 | AMG; b-lactams; TMP-SUL | | OmpK36-0% | |
| OND410A | E | urine | 76 | b-lactams; COL | | - | |
| OND411 | E | rectal swab | 1229 | AMG; b-lactams; TMP-SUL; TET | | OmpK35-37%;OmpK36-31% | |
| OND415 | E | urine | 2800 | AMG; b-lactams; TMP-SUL; TET | | - | |
| OND418 | E | urine | 35 | TET | | - | |
| OND426 | E | blood | 45 | b-lactams; TET; COL | | - | |
| OND447 | E | ascitic fluid | 147 | AMG; b-lactams; TMP-SUL; TET; RIF; MAC; CHL; COL | | OmpK35-59%;OmpK36-34% | |

**Table S2** – Cross-resistance assessment between OMKP-3 and other common antibiotics from different classes (aminoglycosides, fluoroquinolones, cephalosporins, carbapenems, penicillins and miscellaneous) by disk diffusion method (halo diameter determination) for wild-type (WT) and OMKP-3-resistant *K. pneumoniae OND234*

| Antibiotic class | Antibiotic tested | WT (halo diameter, mm) | Fenotype | Resistant mutant (halo diameter, mm) | Fenotype |
|---|---|---|---|---|---|
| aminoglycoside | gentamicin | 13 | R | 13 | R |
| fluoroquinolones | ciprofloxacin | 27 | S | 29 | S |
| cephalosporin | cefoxitin | 15 | R | 15 | R |
| cephalosporin | cefotaxime | 12 | R | 14 | R |
| cephalosporin | ceftazidime | 0 | R | 0 | R |
| cephalosporin | ceftazidime-avibactam | 12 | R | 12 | R |
| carbapenem | imipenem | 19 | I | 21 | I |
| penicillim | temocillin | 10 | R | 10 | R |
| penicillin | ticarcillin | 0 | R | 0 | R |
| penicillin | amoxicillin-clavulanic acid | 0 | R | 0 | R |
| penicillin | piperacillin-tazobactam | 6 | R | 6 | R |
| miscellaneous | sulfamethoxazole | 0 | R | 0 | R |
