## Supplementary data for "*In Vitro* Activity of a Novel Metal-Based Antimicrobial against Multidrug-Resistant *Klebsiella pneumoniae*"

**Supplementary material (figures):**

**Figure S1** – OMKP-3 chemical structure

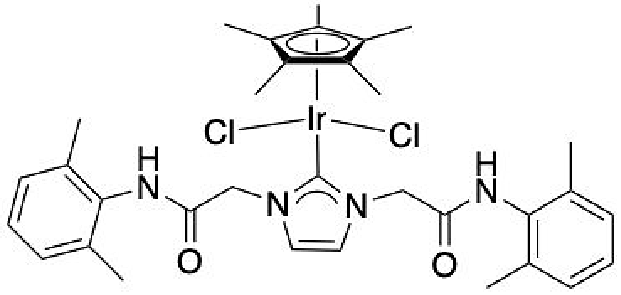

**Figure S2** – OMKP-3 antibiofilm ability against MDR *Klebsiella pneumoniae* strains S10p9A and OND234: (A) OMKP-3 biofilm prevention ability (%); (B) OMKP-3 mature biofilm inhibition (%).

A

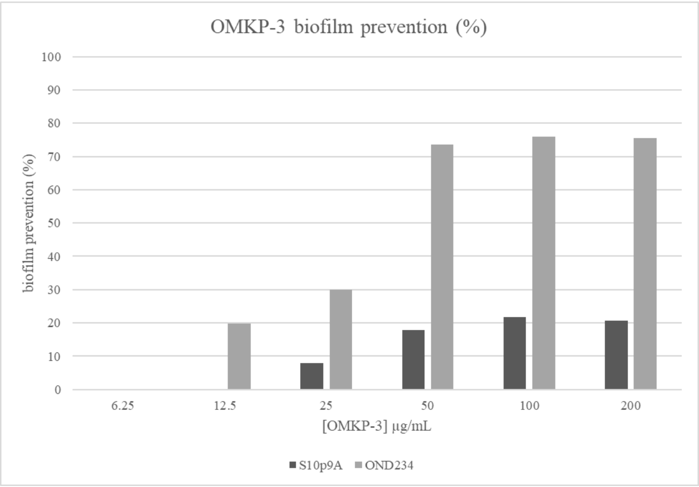


B

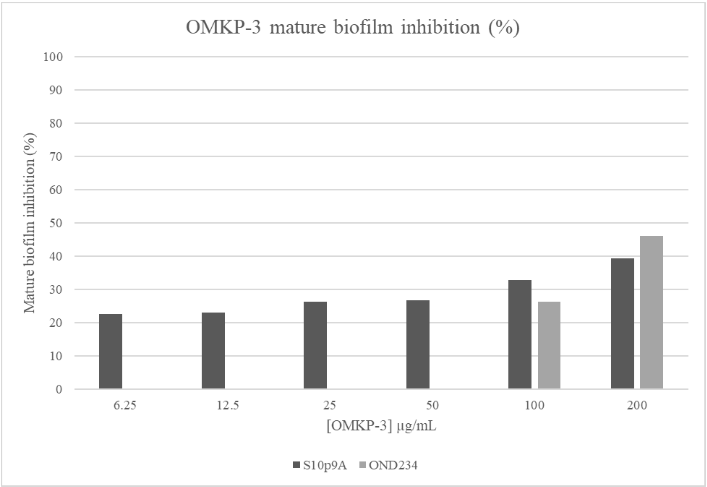

**Figure S3** – Porin (Ompk35 and Ompk36) gene truncation prevalence (%) among MDR *Klebsiella pneumoniae* clinical isolates according to their OMKP-3 minimum inhibitory concentration (MIC) (12.5 µg/mL or 50 µg/mL).

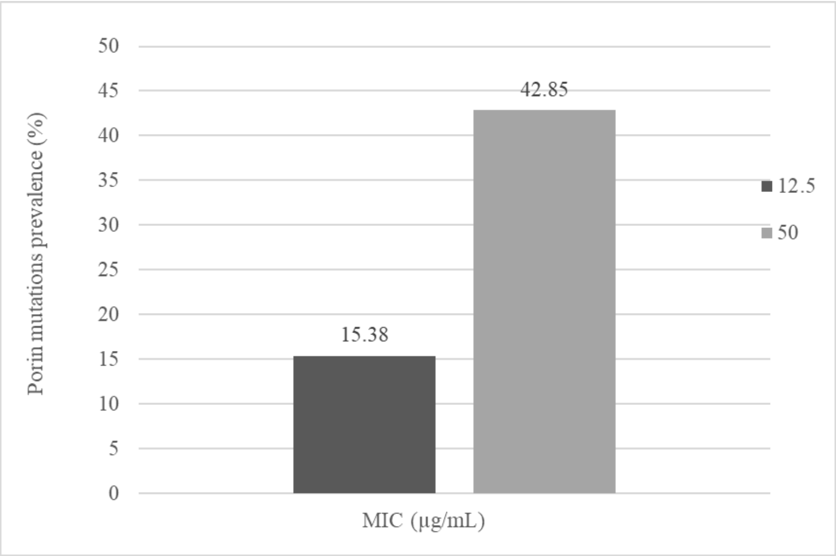

**Figure S4** – Live/Dead bacterial staining images acquired from fluorescence microscopy for *Klebsiella pneumoniae* OND234 strain after OMKP-3 treatment (MIC, 20 minutes). Positive control: boiled bacterial cells (95ºC); negative control: cells treated with M9 minimal media + 0.2% DMSO.

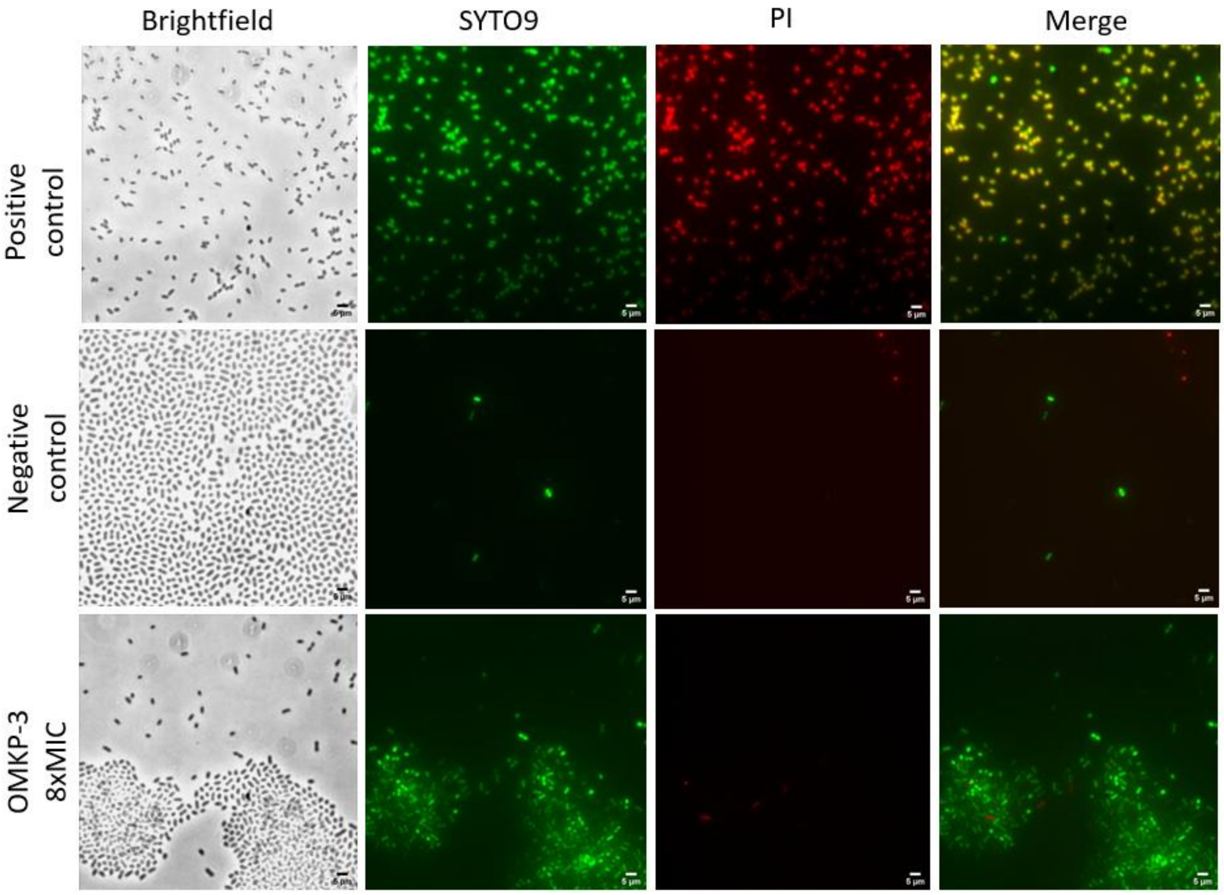
